# Comparative genomics of *Rickettsiella* bacteria reveal variable metabolic pathways potentially involved in symbiotic interactions with arthropods

**DOI:** 10.1101/2024.12.03.626579

**Authors:** Anna M. Floriano, Adil El-Filali, Julien Amoros, Marie Buysse, Hélène Jourdan-Pineau, Hein Sprong, Robert Kohl, Ron P. Dirks, Peter Schaap, Jasper Koehorst, Bart Nijsse, Didier Bouchon, Vincent Daubin, Fabrice Vavre, Olivier Duron

**Author notes:** Corresponding authors: Anna M. Floriano, Olivier Duron.

## Abstract

Members of the *Rickettsiella* genus (order: *Legionellales*) are emerging as widespread bacteria associated with insects, arachnids, and crustaceans. While some *Rickettsiella* strains are highly virulent pathogens, others are maternally inherited endosymbionts that manipulate arthropod phenotypes, including the induction of defensive symbiosis and cytoplasmic incompatibility. However, the genomic diversity of *Rickettsiella* remains largely unexplored, and their genetic potential to induce complex phenotypes in arthropods is only partially understood. In this study, we sequenced five new *Rickettsiella* genomes isolated from three tick species. Through comparative genomics, we observed that *Rickettsiella* members share similar metabolic capabilities, and collectively lack virulence genes from pathogenic Legionellales. Additional analysis of *Rickettsiella* genomes revealed significant variability in metabolic properties related to endosymbiosis. Specifically, their capacity to biosynthesize certain B vitamins and heme varies, suggesting a functional role of some *Rickettsiella* strains in the nutrition of their arthropod hosts. Some *Rickettsiella* genomes harbour homologs of *Wolbachia* cif genes, the cause of *Wolbachia*-induced cytoplasmic incompatibility,, suggesting that *Rickettsiella* may use a similar molecular mechanism to manipulate the reproduction of their arthropod hosts. Phylogenomics further revealed that tick-borne *Rickettsiella* exhibit distinct evolutionary origins within the genus, indicating that *Rickettsiella* have undergone repeated horizontal transfers between ticks and other arthropods.

## Introduction

Bacteria of the genus *Rickettsiella* (order: *Legionellales*, family: *Coxiellaceae*) are emerging as both widespread and genetically diverse in arthropods [1–7]. Historically, *Rickettsiella* have been identified as contagious entomopathogens infecting a broad range of insects, but various pathotypes have also been reported in other arthropods, including arachnids and crustaceans [1–3, 8–11]. In infected arthropods, *Rickettsiella*-induced diseases often replace body cavity tissue with a white iridescent liquid, leading to the death of hosts [9–11], which has spurred research into using pathogenic *Rickettsiella* as a biocidal agent [12],[13]. However, further surveys have revealed that non-pathogenic *Rickettsiella* are common in field populations of arthropods [14–18]. Most of these non-pathogenic *Rickettsiella* are predominantly maternally transmitted, and have a variety of repercussions on the arthropod’s phenotypes, ranging from providing fitness advantages to manipulating reproduction [16, 19–21]. In aphids, *Rickettsiella viridis* can alter body colour, reducing attractiveness to predators and parasitoids [16, 21], or directly protect from entomopathogenic fungi, significantly lowering aphid mortality [16, 19]. In the poultry red mite *Dermanyssus gallinae*, *Rickettsiella* may be an essential nutritional endosymbiont providing B vitamins that are lacking in the blood diet of its host [22]. In the agricultural spider *Mermessus fradeorum*, a *Rickettsiella* endosymbiont induces cytoplasmic incompatibility (CI), that results in embryonic mortality in the offspring of matings between infected males and uninfected females [20]. This reproductive manipulation reduces the production of uninfected progeny and increases the proportion of infected females (*i.e.*, the transmitting sex), thereby promoting the spread of CI-inducing bacteria within the host population [23]. Overall, members of the genus *Rickettsiella* have evolved complex lifestyles that can significantly impact the ecology and evolution of their arthropod hosts [24].

*Rickettsiella* is remarkably common in the two main tick families, Ixodidae (hard ticks) and Argasidae (soft ticks), with at least 15% of species infected and prevalence ranging from 5% to 100% at the population level [4, 5, 25–31]. *Rickettsiella* also exhibit significant genetic diversity both within and between tick species [4, 5, 25–31]. Indeed, genetically distinct strains of *Rickettsiella* were detected in populations of the castor bean tick *Ixodes ricinus* and the polar seabird tick *Ixodes uriae* [25, 26]. The *Rickettsiella*-induced phenotypes in ticks have not been clearly identified yet, but a *Rickettsiella* endosymbiont was observed in a parthenogenetic laboratory colony of the wood tick *Ixodes woodi* [30, 31]. Both males and females are normally present in this tick species, hence *Rickettsiella* might manipulate the tick reproduction to induce asexuality [30]. A bacterium was identified in *I. ricinus*, and further in the blacklegged tick *Ixodes scapularis* [32] as a new genus and species, *Diplorickettsia massiliensis* [33], but it was further synonymised as *Rickettsiella massiliensis* [34] since the bacterium is nested within the current *Rickettsiella* genus [31]. *Rickettsiella massiliensis* has been proposed as a potential agent of opportunistic infections in humans [35], but no clear clinical cases have been reported to date.

*Rickettsiella* is currently among the lesser-known tick-borne bacteria [24], whereas other Legionellales, particularly those that infect humans, are better understood. The order Legionellales includes two families, Legionellaceae and Coxiellaceae, with several clinically significant representatives, such as *Legionella pneumophila*, the causative agent of Legionnaires’ disease, and *Coxiella burnetii*, responsible for Q fever, both of which have been the focus of extensive genomic studies [24, 36]. These pathogenic Legionellales have evolved specific mechanisms to infect mammalian alveolar macrophages, such as a *Dot*/*Icm* type IV secretion system to translocate effectors and inhibit host cell apoptosis [37, 38]. However, not all Legionellales are pathogens: Some are B vitamin provisioning endosymbionts essential for the nutrition of arthropods with an obligate hematophagous lifestyle [22, 24, 36, 39, 40], including *Coxiella*-like endosymbionts in ticks [41–43] and *Legionella polyplacis* in rat lice [44]. Blood is nutritionally unbalanced, and these obligate hematophagous parasites have evolved narrow associations with B vitamin provisioning symbionts, as they themselves cannot synthesise the essential cofactors and vitamins that are lacking in their diet [39, 40]. Additionally, *Coxiella*-like endosymbionts produce chorismate, a tryptophan precursor that regulates serotonin biosynthesis, actively promoting tick feeding activity and blood intake, and modifying behaviour [45].

Compared to other Legionellales, few *Rickettsiella* genomes have been sequenced. They have circular 1.3-1.9 Mb genomes, smaller than the genomes of other *Legionellales* such as the pathogens *L. pneumophila* (3.4 Mb) and *C. burnetii* (2.0 Mb), and the members of the genera *Aquicella* (2.5 Mb) and *Berkiella* (3 - 3.6 Mb), associated with amoebae living in aquatic environments [24, 34, 46]. While *Rickettsiella* genomes are known to encode 1,400-2,200 protein-coding genes [2, 22, 47, 48], no comparative studies have been conducted. So far, the genomes have been used in phylogenomic studies [34, 42], and a few investigations have been conducted focusing on single strains. Indeed, the *Rickettsiella* genome of the poultry red mite *D. gallinae* contains more than 300 pseudogenized protein-coding sequences, but has retained several B vitamin biosynthesis pathways, suggesting the importance of these pathways in the evolution of a nutritional endosymbiosis [22]. Other examples include the genomes of *R. grylli,* infecting isopods, and *R. viridis,* infecting aphids, which both contain a complete *Dot*/*Icm* type IV secretion system [47, 48]. Previous studies have pointed at R. viridis as being a mutualist [14] and at the Rickettsiella isolated from pillbugs as being present in symptomatic (*R. grylli*, [118]), but also in asymptomatic individuals (potentially a different strain, and in coinfection with *R. grylli* [11]). However, many phenotypically characterised *Rickettsiella* strains, as the one inducing CI in spiders [20], have never been sequenced. Among tick-borne *Rickettsiella*, only *R. massiliensis* has had its genome sequenced [2].

In this study, we present five new genomes of *Rickettsiella* naturally infecting ticks, retrace their evolutionary history, characterise their genomic content, and infer their metabolic capacities. To this aim, we sequenced five microbial metagenomes, including three from *I. ricinus*, one from *Ornithodoros erraticus* and one from *Ornithodoros phacochoerus*. We then compared these genomes with published genomes of *Rickettsiella* and representative *Legionellales*, either pathogenic (*L. pneumophila, C. burnetii*) or symbiotic (*Coxiella-*like endosymbionts of ticks), to assess gene content and metabolic similarities. We further used these genomic analyses to infer how *Rickettsiella* can affect the phenotype of their arthropod hosts.

## Methods

### Sample collection, DNA extraction and sequencing

Specimens of three tick species, *I. ricinus*, *O. erraticus* and *O. phacochoerus*, were collected in this study (see Table 1 for details). No data on sex ratio of these tick species were collected in this article nor in previous studies. Different methods for DNA extraction and sequencing were applied depending on tick species. In all cases, total DNA (tDNA) was extracted from the obtained pellet using the DNeasy Blood and Tissue Kit (Qiagen). The obtained tDNA was quantified on Qubit using the dsDNA high-sensitivity kit (Invitrogen). Finally, a shotgun metagenomics approach was applied, obtaining reads belonging to hosts, *Rickettsiella* strains, and any other microbial organisms infecting the same ticks. For the metagenomes Iric, Ird1, and Ird6, DNA extracted from whole tick bodies was first sequenced using PromethION flow cells (FLO-PRO002) with the Ligation Sequencing Kit (SQK-LSK110) from Oxford Nanopore Technologies, and simultaneously sequenced on the NovaSeq 6000 platform with the Nextera DNA Flex Library Prep Kit (Illumina). For the metagenomes Oerr and Opha, endosymbionts DNA samples, enriched as in [41, 51], were sequenced using HiSeq2000 technology using the TruSeq Nano DNA library construction and HiSeq SBS v3 kits (Illumina).

**Table 1:** Collection and processing of samples. * The date 2016 refers to the collection of the ticks in Portugal from which the laboratory colony derived.

| Tick species | Strain | Collection date | Location | Environmental context | Sampling method | Number of ticks | Enrichment |
| --- | --- | --- | --- | --- | --- | --- | --- |
| *Ixodes ricinus* | Iric | 2020 | Netherlands: De Dorschkamp | Deciduous forest | Dragging | 12 (male) | - |
|  | Ird1 | 2021 | Netherlands: De Amsterdamse Waterleiding Duinen | Coastal sand dune area | Dragging | 1 (female) | - |
|  | Ird6 | 2021 | Netherlands: De Amsterdamse Waterleiding Duinen | Coastal sand dune area | Dragging | 1 (female) | - |
| *Ornithodoros erraticus* | Oerr | 2016* | Laboratory colony | Laboratory colony | - | 3 (female) | As described in [41, 51] |
| *Ornithodoros phacochoerus* | Opha | 2021 | Mozambique: Coutada 9 Game Reserve | *Phacochoerus africanus* burrows | Dragging | 4 (female) | As described in [41, 51] |

### Genome assembly

Different methods were used for the Ornithodoros and Ixodes samples. For the Oerr and Opha metagenomes, the reads were subjected to a modified version of the Blobology pipeline [52], available on github. The modified version allows an initial filtering of the host reads using the genome of *I. scapularis* (GCA_016920785.2) as reference, to ease the toll on computing resources, and includes the use of Python3, SPAdes v.3.15.4, bowtie2 v.2.1.0, and blast v.2.12.0. Finally, the resulting genome assemblies were refined with Bandage v.0.8.1 [53], by manual removal of contigs unrelated to the assembly graph with no BLAST hits against the *R. viridis* genome as reference (GCF_003966755.1).

For the metagenomes Iric, Ird1 and Ird6, Minimap2 v.2.24 was used to map all nanopore reads against five known *Rickettsiella* genome sequences (GCF_000168295.1_ASM16829v1, GCF_000257395.1_ASM25739v1, GCF_001881495.1_ASM188149v1, NZ_AP018005.1 and NZ_CP079094.1). Reads that mapped against the *Rickettsiella* reference genomes were collected using Samtools v.1.15.1 and then *de novo* assembled using Flye v.2.9.2_b1786 (error rate 0.03), which resulted in 43 contigs with sizes ranging from 0.5 kb to 1.81 Mb. Medaka v.1.7.2 was used to polish the contigs based on the filtered *Rickettsiella* nanopore reads.

The completeness of retrieved Metagenome-Assembled Genomes (MAGs hereafter) was estimated using miComplete v.1.1.1 [54] (--hmms Bact105). The completeness estimation was performed on a dataset including 12 representative species of *Legionellales* available on NCBI and using the larger genome of the aquatic *Legionellales* “*Candidatus* Berkiella aquae” (GCF_001431295.2) as reference (Supplementary Table 1). This dataset of organisms was used in several analyses in our study and we will therefore refer to it as “Dataset R”. The creation of this dataset was completed on February 1^st^, 2024. Three *R. isopodorum* and one *R. grylli* genome sequences were excluded from the dataset as they do not show significant differences in terms of sequence similarity (ANI higher than 98, calculated with the ANI calculator implemented in EZBioCloud [55]) and gene content (no strain specific ortholog clusters) when compared to their reference genomes. A summary of the organisms included in Dataset R and their hosts is presented in Table 1, while more information about the dataset, including the average nucleotide identity among genomes, can be found in Supplementary Table 1.

**Table 2:**
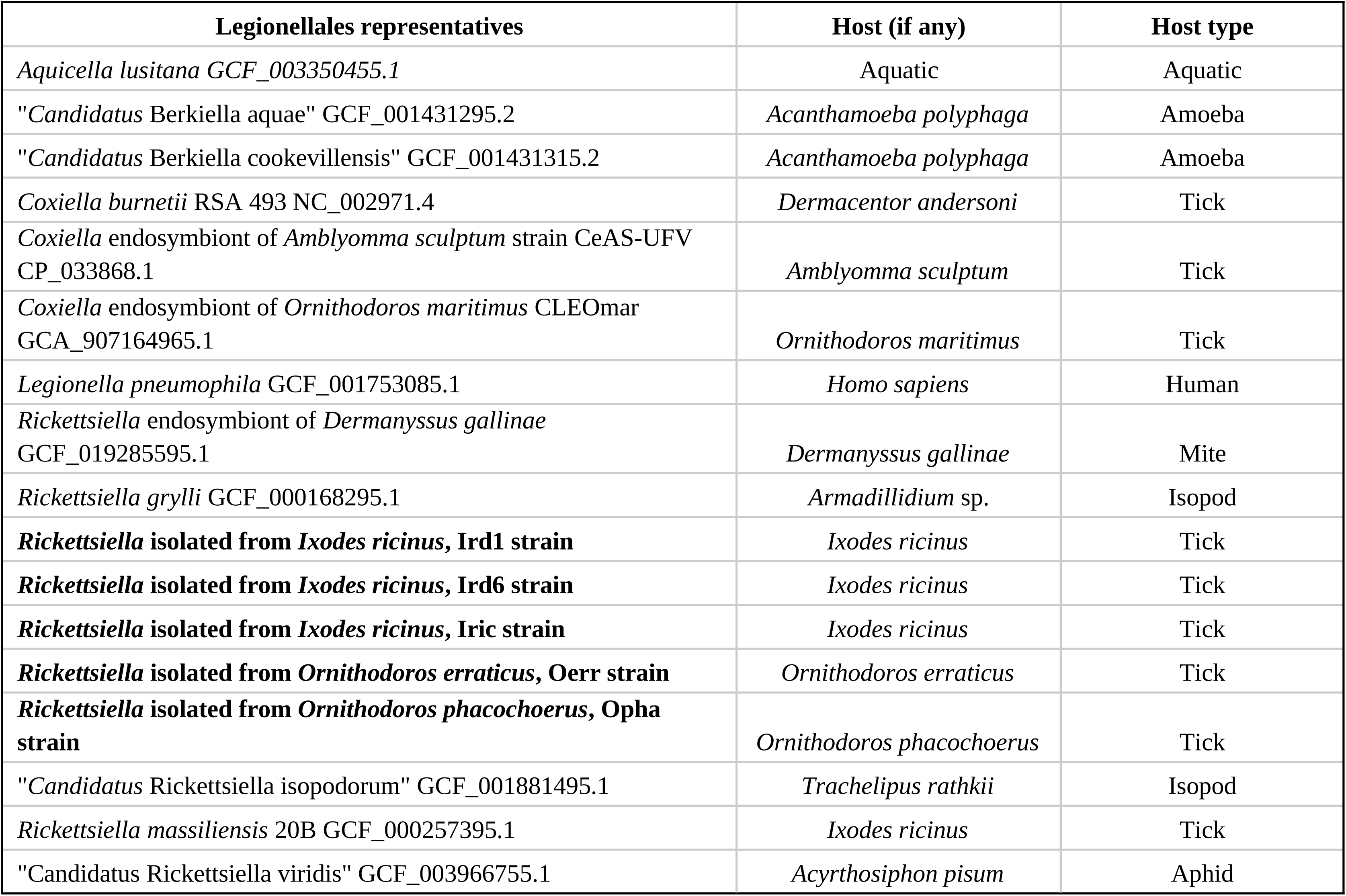
Summary of Dataset R, used for phylogenomics and comparative genomics analyses. The Legionellales representatives’ host scientific and generic names are reported. Strains sequenced in this study are reported in bold.

| **Legionellales representatives** | **Host (if any)** | **Host type** |
| --- | --- | --- |
| *Aquicella lusitana* GCF_003350455.1 | Aquatic | Aquatic |
| "*Candidatus Berkiella aquae*" GCF_001431295.2 | *Acanthamoeba polyphaga* | Amoeba |
| "*Candidatus Berkiella cookevillensis*" GCF_001431315.2 | *Acanthamoeba polyphaga* | Amoeba |
| *Coxiella burnetii* RSA 493 NC_002971.4 | *Dermacentor andersoni* | Tick |
| *Coxiella* endosymbiont of *Amblyomma sculptum* strain CeAS-UFV CP_033868.1 | *Amblyomma sculptum* | Tick |
| *Coxiella* endosymbiont of *Ornithodoros maritimus* CLEOmar GCA_907164965.1 | *Ornithodoros maritimus* | Tick |
| *Legionella pneumophila* GCF_001753085.1 | *Homo sapiens* | Human |
| *Rickettsiella* endosymbiont of *Dermanyssus gallinae* GCF_019285595.1 | *Dermanyssus gallinae* | Mite |
| *Rickettsiella grylli* GCF_000168295.1 | *Armadillidium* sp. | Isopod |
| ***Rickettsiella* isolated from *Ixodes ricinus*, Ird1 strain** | *Ixodes ricinus* | Tick |
| ***Rickettsiella* isolated from *Ixodes ricinus*, Ird6 strain** | *Ixodes ricinus* | Tick |
| ***Rickettsiella* isolated from *Ixodes ricinus*, Iric strain** | *Ixodes ricinus* | Tick |
| ***Rickettsiella* isolated from *Ornithodoros erraticus*, Oerr strain** | *Ornithodoros erraticus* | Tick |
| ***Rickettsiella* isolated from *Ornithodoros phacochorus*, Opha strain** | *Ornithodoros phacochorus* | Tick |
| "*Candidatus Rickettsiella isopodorum*" GCF_001881495.1 | *Trachelipus rathkii* | Isopod |
| *Rickettsiella massiliensis* 20B GCF_000257395.1 | *Ixodes ricinus* | Tick |
| "*Candidatus Rickettsiella viridis*" GCF_003966755.1 | *Acyrtosiphon pisum* | Aphid |

### Genome annotation

Genomes were annotated with Prokka v.1.14.6 [56] using the “compliant” mode and default parameters. The resulting annotations in GenBank format, together with the other 12 annotated *Legionellales* of Dataset R downloaded from NCBI, were subjected to pseudogene analysis with PseudoFinder v.1.1.0 [57] with default parameters. The counts of intact and pseudo-genes have been graphically represented with an in-house R script available on github. KEGG orthology assignments of intact genes and pseudogenes of each genome were predicted using BlastKoala v.3.0 [58]. The outputs were then merged using in-house scripts available on github. Finally, we screened for mobile elements with MobileElementFinder v.1.1.2 [59] using the database MGEdb v.1.1.1 and default parameters, for prophages with PHASTEST v.3.0 [66], and for defense systems (CRISPR/CAS) with CRISPRCasFinder 1.1.2 [116] as in [117].

### Phylogenomics

The phylogenetic relationships between genomes in Dataset R were inferred through a phylogenomic approach on single copy orthologous amino acid sequences (of intact genes, as predicted by PseudoFinder v.1.1.0 [57], alignment length 27116 aa) present in all organisms using *Methylococcus mesophilus* strain 16-5 (GCF_026247885.1, intact genes predicted the same way) as the outgroup. The pipeline is available on GitHub and includes the use of the tools OrthoFinder v.2.5. [60], SeaView v.5.0.5 [61], ModelTest-NG v0.2.0 [62], RAxML (raxmlHPC-PTHREADS v.8.2.12, [63], 1000 bootstraps), and in-house scripts in Python3, bash, and R. All of the softwares have been run in default settings, except for RAxML, where the model was set to LG+I+G4 (PROTGAMMAILG), chosen according to the Akaike criterion with ModelTest-NG v0.2.0.

### Overall comparative genomics

In order to obtain a description of common and strain-specific genomic features and metabolic capabilities of *Rickettsiella*, we analysed the clusters of orthologous intact genes detected for Dataset R, regardless of phylogenetic relationships. The OrthoFinder output table from which the single-copy orthologs used in the phylogenomic inference (Orthogroups.GeneCount.tsv, Supplementary Table 2) was analysed for intersections using the R package UpSetR [67], and the results were plotted using the same tool. All the commands are available on github.

Particular attention was paid to clusters uniquely shared by (i) all *Rickettsiella* species and strains, (ii) *Rickettsiella* strains isolated from hematophagous arthropods, for traces of adaptation to host blood feeding (iii) *Rickettsiella* strains isolated from phytophagous/detritivorous hosts, for traces of adaptation to other diets (iv) *Rickettsiella* isolated from the same host, for adaptation to hosts/environments and (v) *Rickettsiella* and *Coxiella* isolated from ticks, for traces of adaptation to tick hosts. We also investigated gene orthology in respect to the phylogenetic clusters that appeared well-defined in the phylogenetic trees. Orthology was assessed using OrthoFinder on all translated proteins predicted in genomes by Prokka, as well as on translated sequences of intact genes and pseudogenes as previously predicted with PseudoFinder, in order to obtain more information about the evolution of the strains. The results were graphically represented as Venn diagrams using the R package VennDiagram [68], and the KEGG and Prokka annotations of the clusters were obtained (commands available on GitHub).

### Comparison of functions of interest

We also analysed specific pathways that could be involved in nutritional symbioses with hematophagous arthropods (B vitamins, chorismate and heme metabolism [45, 51, 69, 70]), using subsets of Supplementary Table 3 to obtain graphical representation of the gene presence/absence/pseudogenization along the phylogenomic tree with an in-house R script available on github. The analysis showed differences in some of the pathways. In particular, the thiamine pathway caught our attention: the strain Iric appears to have a set of genes that are absent in other Rickettsiella strains but shared with other Legionellales representatives. We thus further investigated these genes (Supplementary Table 3) by retrieving additional sequences from NCBI, identified via blastp, and aligned the amino acid sequences with MUSCLE v3.8.1551. Finally, we inferred their respective phylogenies with RAxML (100 bootstraps).

We further checked for the presence of virulence factors known in *Legionellales* [37, 38]. Protein sequences of virulence factors annotated in *Legionella longbeachae* NSW150 (NC_013861), *L. pneumophila* str. Corby (NC_009494) and *L. pneumophila* subsp. *pneumophila* str. Philadelphia 1 (NC_002942) were downloaded from VFDB [71] and used as queries in a blastp search against the translated sequences of intact genes and pseudogenes of genomes from Dataset R.

Finally, we also looked for the presence of CI genes. The CI phenotype is encoded by a pair of syntenic genes, *cifA* and *cifB*, whose direct genotype-phenotype association has been demonstrated in *Wolbachia* [72, 73]. *cifA* and *cifB* possess homologs that have been categorised into five phylogenetic clusters [74–77], although type V has recently been split into type V-to-X [77]. To determine the presence of *cif* gene homologs in the *Rickettsiella* genomes, we used Orthofinder to search for specific orthologs of the *cif* gene groups identified in a previous study [74]. These searches were conducted on a database of CDSs from Dataset R, as well as from published *Wolbachia,* known to induce CI, and *Rickettsia* genomes (Supplementary Table 4). Protein domains were predicted for the *cifA* and *cifB* homologs detected using the HHpred webserver (https://toolkit.tuebingen.mpg.de/#/tools/hhpred/, [78]) with default parameters [79], and using the following databases: SCOPe70 v.2.08, Pfam v.36.0, SMART v.6.0, and COG/KOG v.1.0. The sequences of each detected domain were retrieved and manually aligned using Clustal Omega [80], independently for each *cifA* and *cifB* homologs, to complete the domain detection of some false negatives generated by HHpred. Visualisation of protein domains within the *cif* genes was performed using RStudio v4.3.1 with the ggplot2 v.3.4.4 [81], cowplot v1.1.1 [82], and gridExtra v.2.3 [83] packages. A maximum likelihood phylogeny of *cifA* and *cifB* pair genes has been inferred on the basis of three conserved domains (RNA-binding-like, AAA-ATPases-like and PD-(D/E)XK nucleases) independently aligned and concatenated. The phylogenetic reconstruction was performed using the best-fitting substitution model, here JTT+G, as identified by ModelTest-NG, based on the Akaike Information Criterion.

## Results

### Rickettsiella metagenomes

We sequenced and assembled three microbial metagenomes from *I. ricinus* (Ird1, Ird6 and Iric), one from *O. phacochoerus* (Opha) and one from *O. erraticus* (Oerr). We obtained an average of 71.86 Gb of paired-end reads (from 2.2 gb to 192 bp) of 150 nt in length for each sample (see Supplementary Table 1 for details). For each metagenome, we extracted the reads belonging to the target organisms and obtained the sequences of five new *Rickettsiella* MAGs (see Supplementary Table 1 for details). They are all of similar size (∼1.7 Mb), with the exception of *Rickettsiella* Oerr MAG (∼1.3 Mb), and most of them are estimated to be complete. However, the estimation attributes to *Rickettsiella* strain Oerr MAG lower completeness scores (0.9048; weighted completeness 0.8109) among the MAGs sequenced in this study. Lower scores are expected for endosymbiotic, host-restricted bacteria, whose genomes undergo erosion [84–87], and observed in *Rickettsiella* relatives, as the *Coxiella*-like endosymbionts of ticks. Thus, the completeness score of *Rickettsiella* strain Oerr MAG reflects more likely genomic erosion than poor assembly, albeit the number of contigs included in this MAG reaches 99. However, it is worth noting that other genomes, previously published and considered to be complete, are assembled in a higher/comparable number of contigs (*Coxiella* CLEOmar:, *Aquicella lusitana*) or present a lower set of markers (*Coxiella* CeAS-UFV, 93 out of 105 markers used in the analysis, two less than *Rickettsiella* Oerr). The GC content varies between 36% and 39%, values also consistent with those from endosymbiotic bacteria. Metagenomes also contain reads of other bacterial endosymbionts as *Midichloria* and *Francisella*-like endosymbiont for *I. ricinus* and *O. phacochoerus*, respectively (data not shown). These endosymbionts were not further considered in this study. No other endosymbiont than *Rickettsiella* was detected for *O. erraticus*.

The annotation of the five *Rickettsiella* MAGs revealed a somewhat similar number of functional genes, ranging from 1,268 to 1,473 (Supplementary Table 1). Including all *Rickettsiella* genomes shows that the number of functional genes is between 1,189 for *R. isopodorum* and 1,707 for the *Rickettsiella* endosymbiont of *D. gallinae* (Supplementary Table 1, Supplementary Figure 1). *Rickettsiella* genomes display similar levels of pseudogenization to other *Legionellales*, though some variation is observed. For instance, higher numbers of pseudogenes were predicted for the *Rickettsiella* MAGs Iric and Oerr (340 and 237 respectively), *R. massiliensis* 20B (508) and a few other *Legionellales*, including *Coxiella*-like endosymbiont of *Ornithodoros maritimus* strain CLEOmar (778). In contrast, lower numbers of pseudogenes were predicted in other *Legionellales*, such as the *Coxiella*-like endosymbiont of *Amblyomma sculptum* strain CeAS-UFV (31) (Supplementary Table 1, Supplementary Figure 1). Concerning mobile elements, only one was detected in the analysed dataset, and only in *R. grylli* (Supplementary Table 1). Both recently host-restricted bacteria and long-term symbionts generally possess a low number of mobile elements in their genomes, although recent symbionts are usually characterised by a greater number. However, long term symbionts are additionally characterised by fewer pseudogenes and smaller genome sizes [85] than the *Rickettsiella* strains here investigated. Taken together, these features suggest a relatively recent infection of the hosts by *Rickettsiella*. In addition to mobile elements, prophage sequences were detected in the strains Iric and Ird6 and in the Rickettsiella endosymbiont of D. gallinae (Supplementary Table 1), but not in other strains. Furthermore, we investigated defense systems to evaluate the degree of adaptation to an endosymbiotic lifestyle as in *Arsenophonus* [117] (Supplementary Table 1).

Most Average Nucleotide Identities (ANIs) between the five *Rickettsiella* MAGs range from 69.39 to 72.61%, suggesting that they may be distinct species (Supplementary Table 1). A notable exception is the 99.77% ANI between Iric and Ird6. The five *Rickettsiella* MAGs have ANIs ranging from 69.36-to-98.46% with other *Rickettsiella* genomes. Indeed, ANI is 98.46% between Iric/Ird6 and *R. massiliensis*, and 98.26% between Ird1 and *R. isopodorum* (Supplementary Table 1).

### Phylogenomics of *Rickettsiella*

Phylogenomic analysis based on 109 single-copy orthologs showed that *Rickettsiella*, *Coxiella*, *Legionella*, *Aquicella*, and *Berkiella* genera cluster together in a robust monophyletic clade forming the *Legionellales* order (Figure 1). The inner tree topology is also congruent with the known genera of *Legionellales* [24]. Indeed, genomes of *Rickettsiella*, *Coxiella*, and *Berkiella* cluster together according to their assigned genus and form robust monophyletic subclades, with *Aquicella* and *Coxiella* forming sister genera of *Rickettsiella*. Two *Rickettsiella* MAGs (Iric and Ird6) obtained in this study from *I. ricinus*, cluster with *R. massiliensis* sharing a high ANI from 98.5 to 99.8% (node 15, Figure 1, and Supplementary Table 1), suggesting that these three *Rickettsiella* belong to the same species. The three other *Rickettsiella* MAGs obtained in this study cluster with diverse *Rickettsiella* species: Opha with R. viridis, the defensive symbiont of aphid, and the putative nutritional *Rickettsiella* symbiont of poultry red mite (node 13), Ird1 with *R. isopodorum* (node 12) and Ierr with a group formed by *R. grylli*, *R. isopodorum* and Irid6 (node 10, Figure 1). The *Rickettsiella* found in different tick species are thus distantly related and do not form a monophyletic group (Figure 1).

**Figure 1:**
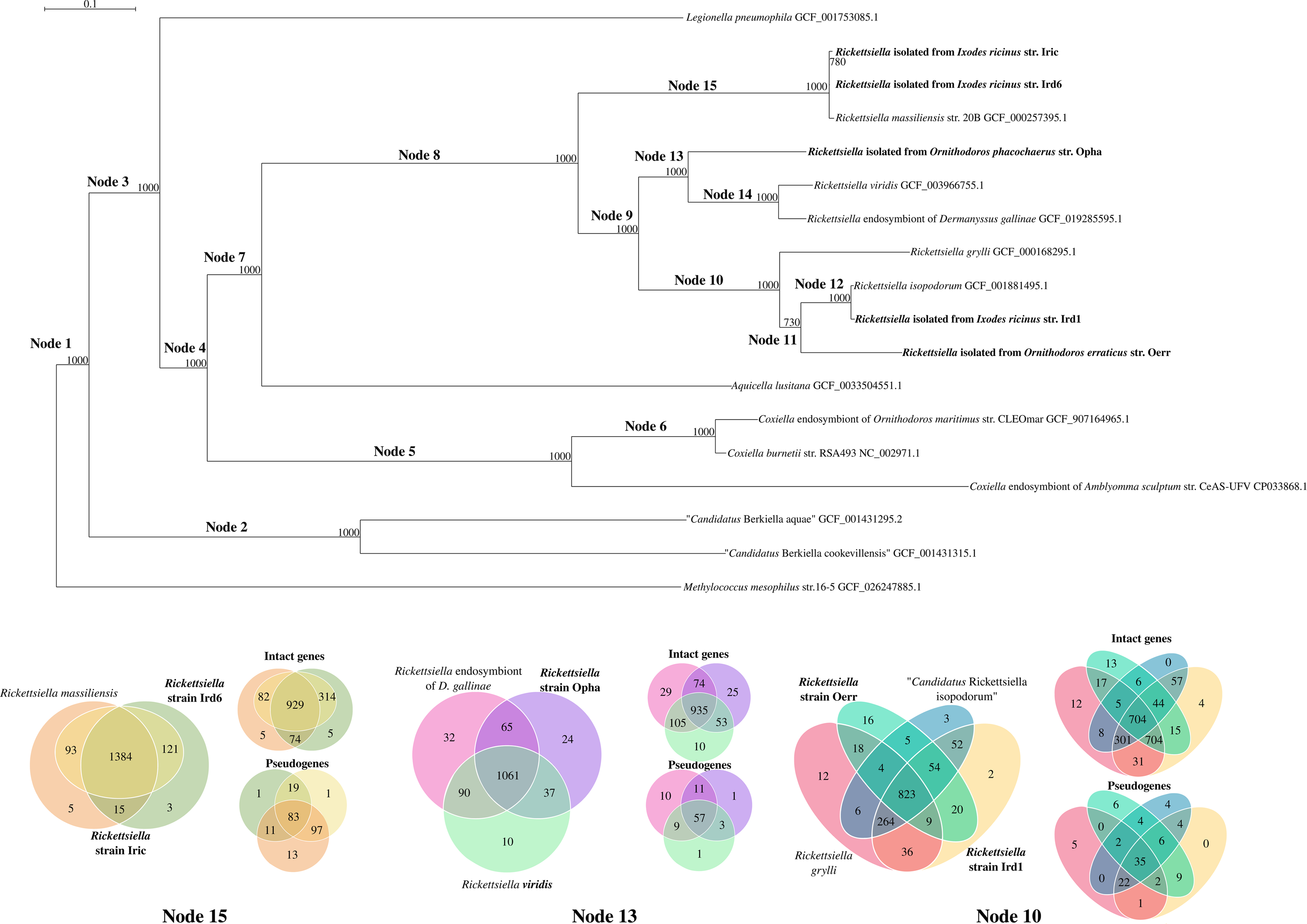
Phylogeny of Legionellales (Dataset R) focusing on the *Rickettsiella* genus inferred with RAxML on 109 single copy orthologous intact genes present in all organisms in the dataset (1000 bootstraps, evolutionary model: LG+I+G4, rapid bootstrap random number seed: 1234, the random number seed for the parsimony inferences: 123), using *Methylococcus capsulatus* as outgroup to root the tree. Only bootstrap supports lower than 98 are shown. The *Rickettsiella* organisms whose genomes were sequenced in this study are indicated in bold. On the right, the presence (green), absence (grey) and pseudogenization (yellow) of genes of specific metabolic pathways annotated with KEGG is represented as a heatmap. *Methylococcus capsulatus* sequences were not annotated. On the bottom, Venn diagrams of clusters of orthologs of all encoded genes, intact genes, and pseudogenes is provided for the specified nodes of the tree (Node 10, Node 13, Node 15).

## Comparative genomics

### Orthology analysis

The analysis of the intact orthologous genes detected for Dataset R (Figure 2, Supplementary Figure 2, Supplementary Table 2) indicates huge variation among Legionellales, with only135 clusters shared by all, most of which are involved in basic bacterial processes. Only 25 clusters of intact genes are shared exclusively by members of the *Rickettsiella* genus, most of which are functionally uncharacterised (Supplementary Table 2). No clusters of intact genes specific to *Rickettsiella* and *Coxiella* strains isolated from hematophagous arthropod hosts were detected. The core *Rickettsiella* genome appears to contain 606 COGs, that drop to 362 when pseudogenes are removed from the analysis; pseudogenes in the *Rickettsiella* core genome (29 clusters of pseudogenized sequences) appear to be mainly associated with transcription and regulation functions (Supplementary Table 5). We then attempted to detect gene families specific to the phylogenetic groups that could be identified on the phylogenomic tree shown in Figure 1.

**Figure 2:**
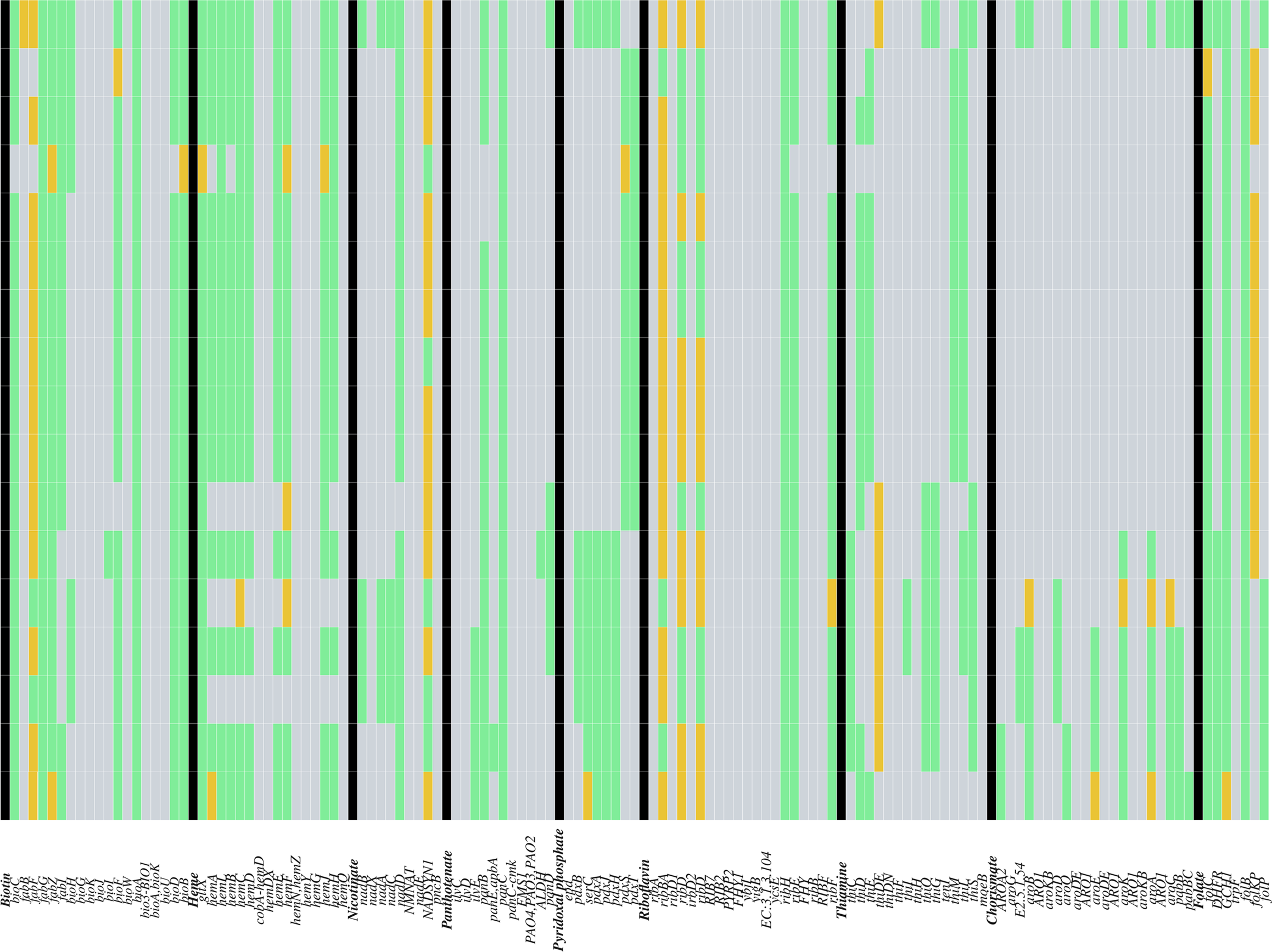
Intersection plot displaying shared and specific clusters of orthologous proteins between *Rickettsiella* and other Legionellales. Intersection plot is based on Dataset R and was produced with the R package UpSetR. In the left side, the horizontal bar plot represents the number of orthogroups present in each organism of the dataset. In the top, the vertical bar plot represents the number of orthogroups that belong to specific intersections of organisms in the dataset. The height of each bar indicates how many orthogroups are in that particular intersection, i.e., shared by the combination of organisms indicated by the intersection matrix below. In the intersection matrix, on the bottom, each row represents an organism and each column represents a possible intersection of these organisms. Filled circles indicate that the organisms in that row are part of the intersection shown in the corresponding column. The lines connecting the filled circles indicate which organisms are included in each specific intersection. This plot is limited to six orthogroups in the intersections, for the full plot of all intersections until one shared orthogroup please see Supplementary Figure 2.

Node 10 encompasses two previously described and potentially pathogenic *Rickettsiella* species isolated from isopods (*R. grylli* [3] and *R. isopodorum* [88]), and two of the newly *Rickettsiella* strains identified in ticks. Notably, these two new strains were isolated from very distant ticks, both in terms of host species and geographic location (Supplementary Table 1). The core genome of these organisms contains 823 COGs, which drop to 704 when pseudogenes are not considered in the analysis (Figure 1). Of the 704 COGs, 198 correspond to uncharacterised proteins, and 27 appear to be involved in secretion of effectors and pumps. Furthermore, five were annotated by Prokka as multidrug resistance proteins. However, the putative pathogen *R. grylli* possesses 22 additional transport proteins compared to the rest of this dataset, and *Rickettsiella* Oerr shows 35 specific uncharacterised proteins (Supplementary Table 5).

Node 13 encompasses the host colour-altering and defensive symbiont *R. viridis* (aphid host, [14]), the *Rickettsiella* endosymbiont of *D. gallinae* (poultry red mite host, [89]), and the newly sequenced *Rickettsiella* str. Oerr isolated from the soft tick *O. phacochoerus*. The core genome of this node contains 935 COGs (1061 when pseudogenes are considered; Figure 1). All the *Rickettsiella* strains in this node appear to maintain a discrete, albeit low, number of unique COGs. Among others, the core intact genes include T4SS components, ATPase subunits, biotin genes, 79 hypothetical proteins, four ankyrin repeat domain-containing proteins, and 14 DUF domain-containing proteins.

Notably, the Opha strain appears to share more COGs with the *Rickettsiella* of *D. gallinae* (65, encoding hypothetical proteins, several ATPases, DUF containing-proteins, and transcription regulators) than with *R.viridis* (37, most of them being potentially involved in adhesion). Interestingly, the core pseudogenes include many response regulators and translocators. Finally, the aerobic respiration control sensor protein ArcB is pseudogenized in the strain Opha only, and the endosymbiont of *D. gallinae* contains 10 more unique pseudogenes, that include regulators and transposases (Supplementary Table 5).

Finally, Node 15 encompasses three *Rickettsiella* strains isolated from the hard tick *I. ricinus*: *R. massiliensis*, *Rickettsiella* Iric and Ird6. The core of 929 COGs (1384 when pseudogenes are included in the analysis) contains mainly hypothetical proteins and house-keeping genes, but also genes involved in energy production. Interestingly, the strain Iric exhibits distinctive COGs exclusively when pseudogenes are analysed, revealing a single cluster associated with amino sugar and nucleotide sugar metabolism (Supplementary Table 5). However, the other strains possess only five unique COGs as well. A relatively high number of intact sequences is shared by any two organisms in these subset of genomes. Most sequences shared by *R. massiliensis* and the strain Iric are annotated as hypothetical proteins, whereas most sequences shared by *R. massiliensis* and the strain Ird6 are housekeeping genes and genes encoding chaperones. The *Rickettsiella* Iric and Ird6 share mostly uncharacterised proteins, but also several dehydrogenases, ATPase synthase subunits A and B, and a queuosine precursor transporter (Supplementary Table 5).

### Metabolic capabilities

The annotation of metabolic pathways (Supplementary Table 3) shows overall similar metabolic capabilities among the members of the *Rickettsiella* genus. Major differences can be identified, however, in specific pathways: the respiration and ATP production appears to be most degraded in strain Iric and *R. massiliensis* 20B, *i.e.*, the two *Rickettsiella* strains with the higher number of predicted pseudogenes that form a monophyletic cluster together with the Ird6 strain.

Furthermore, B vitamins and heme pathways show significant differences, appearing most degraded in *R. massiliensis* 20B, but still relatively conserved in the strain Iric (*e.g.*, biotin, folate). Strikingly, the heme biosynthesis pathway is absent in the strain Oerr, which has also lost most of the thiamine biosynthesis pathway, while retaining a unique segment of this pathway that is not found in other *Rickettsiella* strains sequenced in this study. However, although it is not possible to fully reconstruct the evolution of these sequences, a blast search and inferred phylogenies for each gene shows that they are of Gammaproteobacteria origin (Supplementary Table 3, five best hits shown, and Supplementary Figure 3), often found in *Legionella* spp. and in two additional *Rickettsiella* organisms. This suggests either multiple horizontal transfers from other Gammaproteobacteria or Legionellales, or a loss of the genes in the strains analysed here.

Finally, while *Coxiella*-like endosymbionts can produce chorismate, a tryptophan precursor that regulates serotonin biosynthesis in ticks [44], *Rickettsiella* genomes do not harbour genes for this biosynthetic pathway, suggesting that they have no impact on tick behaviour through this mechanism.

### Identification of virulence factors

Further genomic comparisons showed that abundance of virulence genes varies between Legionellales members (Figure 4, Supplementary Table 6). While large repertories of virulence genes associated with adherence, secretion systems, effectors and exotoxin RtxA are common in *L. pneumophila* genomes, these genes are rare or absent in most other Legionellales as *Coxiella*-like endosymbionts associated with ticks. Indeed, some virulence genes are absent in *Rickettsiella* genomes including the exotoxin RtxA-encoding locus, which is involved in adherence, cytotoxicity, and pore formation in vertebrate cells by *Legionella* species [90]. *Aquicella lusitana* also lacks the exotoxin, suggesting that the loss of this gene might be ancestral to *Rickettsiella* and *Aquicella*. As with the other factors associated with virulence in *Legionella*, *Rickettsiella* genomes conserve those involved in adherence, regulation, and nutritional metabolism (particularly, heme metabolism cofactors [91]). Furthermore, the *Rickettsiella* strains retain the effector delivery systems, but not the secreted virulence-associated effectors identified in *Legionella*, suggesting that this machinery could be exploited by *Rickettsiella* to perform different functions. The same applies to *Aquicella*, *Berkiella* and *C. burnetii*, whereas the two *Coxiella*-like endosymbionts have neither the delivery system nor the genes encoding for effectors. Furthermore, all the investigated *Legionellales*, with the exception of *Legionella* and several *Rickettsiella* strains, do not possess the Type IV A secretion system (TIVASS). The strains that maintain most of this system are *R. viridis* and the tick strains Ird1, Oerr and Opha, while the *Rickettsiella* endosymbiont of *D. gallinae*, *R. grylli* and *R. isopodorum* have only a few genes involved in this machinery. No evidence of horizontal gene transfer was found. Finally, no enhancer-binding protein (Enh) encoding loci were found in *Rickettsiella*, whereas they are found in *Aquicella*, *Berkiella*, *C. burnetii*, and *Legionella*.

### *Rickettsiella* and cytoplasmic incompatibility

We detected homologs of the CI genes *cifA* and *cifB* in the genomes of (i) *Rickettsiella* strain Opha, and (ii) *R. isopodorum* where a CifB-like protein has been previously identified (Figure 3, [92]). Both strains harbour the RNA-binding-like domain in *cifA*, and the AAA-ATPase-like and PD-(D/E)XK nuclease domains in *cifB*, both conserved in all known *cif* genes [74, 93]. Additionally, the *cifB* gene in the strain Opha has a weakly aligned latrotoxin-like fragment, while *R. isopodorum* has a large TcdA/TcdB pore-forming toxin domain, also found in other *cifB* homologs of *Wolbachia* and *Rickettsia*. Furthermore, we also identified a unique *cifB* homolog (without *cifA*) in a *Rickettsiella* sp. genome reconstructed from eDNA isolated from hot springs and volcanic lakes (GCA_037439325.1) and in the distant relative “*Ca.* Aquirickettsiella gammari” (GCA_002290645.2).

**Figure 3:**
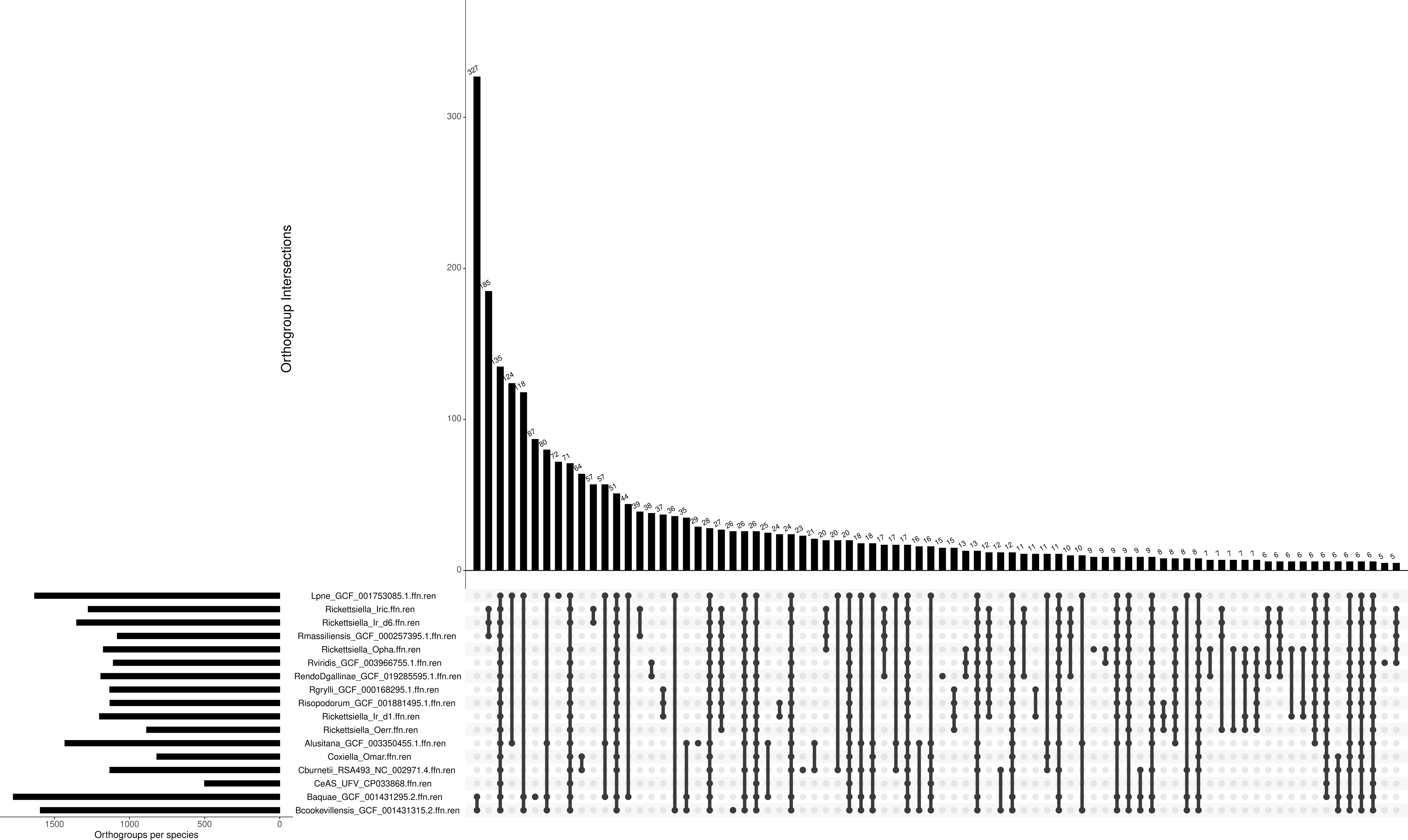
Phylogeny and domain architecture composition of cifA and cifB homologs in *Rickettsiella* and other endosymbionts. The maximum-likelihood tree was reconstructed from an alignment concatenated with RNA-binding-like, AAA-ATPase-like, and PD-(D/E)XK nuclease domain sequences (191 amino acid positions, JTT+G substitution model). The tree is midpoint rooted. The type I-V cif representation, as well as the sizes and domain compositions of the cifA and cifB genes, were drawn relative to those described in Martinez et al. (2021, 2022). The scale bar represents the inferred number of substitutions per site. Bootstrap values were estimated from 1000 replicates. Accession numbers of the *Rickettsiella*, *Wolbachia*, and *Rickettsia* genomes used in this analysis are listed in Supplementary Table S4.

**Figure 4:**
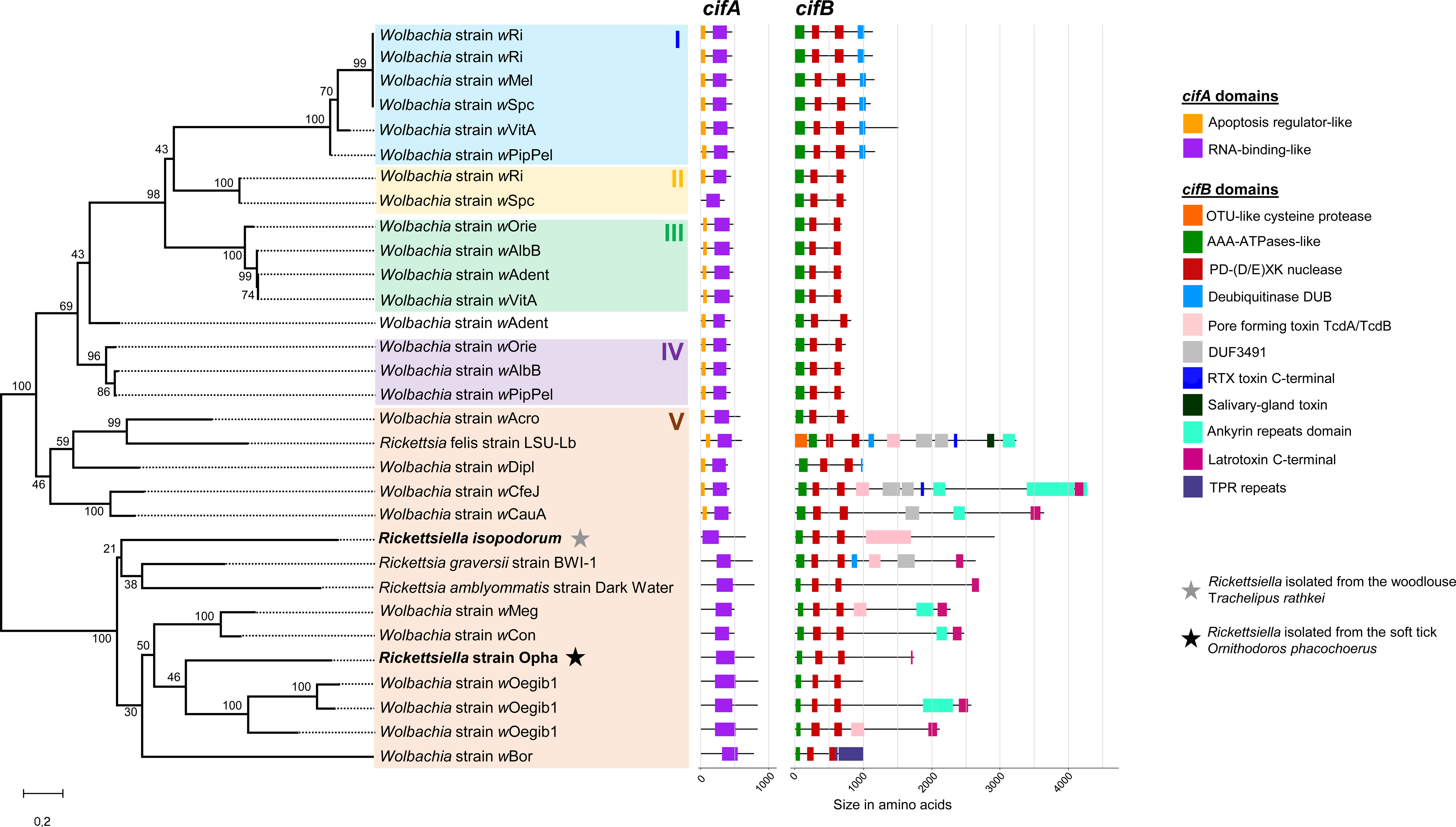
Barplot of the virulence factors present in each organism of our dataset. The virulence factors are grouped by category. The presence is expressed as the percentage of genes present for each category (i.e., when a category includes ten components and ten of them are present in an organism, this is shown as 100%). Commands for the plot production are available on GitHub.

Both the detected *cifA* and *cifB* homologs of *Rickettsiella* cluster within the Type V *cif* paraphyletic group, which is the most diverse in terms of size and domain composition [74, 93]. These genes are located in the chromosomal DNA and without any nearby mobile genetic elements, such as prophages, insertion sequences (IS), or retrotransposons. Both the *cif* genes from *Rickettsiella* strain Opha and *R. isopodorum* group together in a highly supported clade comprising *cif* genes from *Rickettsia* and *Wolbachia*, originating from at least two supergroups (*w*Oegib and *w*Bor from supergroup A, *w*Meg and *w*Con from supergroup B), showing the occurrence of gene transfers between these phylogenetically distinct endosymbionts. However, the long branch lengths and low intra-group bootstrap values create uncertainty regarding the exact position of the *Rickettsiella cif* genes, challenging the reconstruction of their evolutionary history and origin.

## Discussion

We show here that members of the genus *Rickettsiella* share a set of similar metabolic capabilities, as pathways involved in energy production. Most notably, the *Rickettsiella* strains lack most key virulence genes responsible for persistence and replication in vertebrate cells of pathogenic *Legionella* and *Coxiella*. Indeed, while the *Rickettsiella* genomes harbour the virulence genes encoding the *Dot*/*Icm* secretion system, the genes encoding the Legionellales *Dot*/*Icm* secreted effectors and the exotoxin RtxA have been pseudogenized or are completely absent. However, examination of *Rickettsiella* genomes further reveals a greater variability in metabolic properties associated with endosymbiosis: They have variable capacity to biosynthesize *de novo* certain B vitamins (biotin, riboflavin, pyridoxal phosphate, pantothenate) and heme, and some harbour CI genes, suggesting that they are engaged in complex interactions with their respective arthropod hosts. Phylogenomics further confirmed that members of the genus *Rickettsiella* form a robust monophyletic clade of Legionellales associated with arthropods but that tick-borne *Rickettsiella* do not cluster within a specific subclade. Tick-borne *Rickettsiella* rather exhibit distinct evolutionary origins, revealing that *Rickettsiella* have undergone repeated horizontal transfers between ticks and other arthropods, including aphids and woodlice.

The virulence factors presence/absence pattern across Legionellales reveals key past events in the evolutionary history of *Rickettsiella*. Indeed, the absence of major Legionellales virulence factors in all *Rickettsiella* strains, including the putative human pathogen *R. massiliensis* 20B, suggests that these microorganisms are not strongly pathogenic to mammals as *L. pneumophila* and *C. burnetii*. *Aquicella lusitana*, *Rickettsiella*’s closest relative, also lacks the RtxA exotoxin, suggesting that the loss of this gene could be ancestral to Rickettsiella and Aquicella and extend deep into the evolutionary past of the Coxiellaceae. Several *Legionella* virulence factors are conserved in *Rickettsiella* and to some extent in *Aquicella*, though they may have alternative functions. Notably, traits such as adherence, stress response regulation, nutritional metabolism (particularly involving heme cofactors), and the type IV secretion systems are common to both pathogenic and mutualistic bacteria [87,90–97] whereas the *Dot*/*Icm* effector delivery system has been so far exclusively associated with pathogenesis [42]. Indeed, genes encoding the *Dot*/*Icm* system are either absent or pseudogenized in *Coxiella*-like endosymbionts associated with nutritional symbiosis in ticks [42,98]. The conservation of this secretion system in *Rickettsiella* suggests either that this machinery is exploited by *Rickettsiella* to fulfill functions other than virulence by secreting different effectors, or that *Rickettsiella* maintains a degree of virulence but uses different virulence effectors to those of *Legionella*. Alternatively, since the *Rickettsiella Dot*/*Icm* system lacks a few genes compared to *L. pneumophila* and *C. burnetii*, this machinery may have undergone a gradual degradation and they mirrored a global genome decay, as commonly observed in endosymbionts of arthropods [84, 85, 87, 99, 100]. Whatever the scenario, virulence against mammals seems to be gradually lost in *Rickettsiella*, leaving opportunities for the evolution of other interactions with their arthropod hosts.

The genomes of *Rickettsiella* consistently harbour genes involved in narrow interactions with arthropods, although these genes vary between trains and species. The biosynthesis genes of several B vitamins (biotin, riboflavin, and folate) were found intact in most *Rickettsiella* genomes, suggesting that they could supply these nutrients to their arthropod hosts. In this context, *Rickettsiella* may be essential mutualistic partners for hematophagous arthropods, such as ticks and certain mites, which need these essential vitamins for their growth and the completion of their life cycle [39]. Most tick species harbour *Coxiella*-like endosymbionts which provide B vitamins, but tick species lacking *Coxiella*-LE instead host *Francisella*-like endosymbionts [39, 101] or *Midichloria* [102, 103] as we observed in this study in *O. phacocherus* and *I. ricinus*, respectively. These symbionts are also capable of supplying B vitamins to their hosts. Co-infections (*i.e.*, the simultaneous infection of *Rickettsiella* with one or more different microorganisms) have been previously documented in ticks [4, 5, 101], hence, the identification of *Rickettsiella* tick species previously associated with other symbionts is not surprising. However, in *O. erraticus*, we did not observe any other endosymbionts than *Rickettsiella* which suggests that this bacterium is an obligate nutritional partner of this host. Current knowledge, including a study on a related tick species [51], shows that a mutualist is necessary. In addition, certain *Rickettsiella* genomes possess biosynthesis genes for heme, a nutrient which is essential for regulating protein synthesis and cell differentiation in animals. Unlike other eukaryotes, ticks lack most of the genes encoding proteins necessary for heme production and degradation [104–106]. Some *Rickettsiella* endosymbionts could thus potentially complement tick metabolism through the production of this essential cofactor.

Further examinations of *Rickettsiella* genomes also suggest that at least two strains may be reproductive parasites of the soft tick *O. phacochoerus* and the woodlouse *Trachelipus rathkei*. Although no phenotypic data are currently available for these arthropod species, their *Rickettsiella* possess CI-inducing genes with key functional domains, including a RNA-binding-like domain in *cifA*, a the AAA-ATPase-like and PD-(D/E)XK nuclease domains in *cifB*, as observed in all *cif* genes of CI-inducing *Wolbachia* [74, 93]. The observed pattern suggests acquisitions of *cif* genes in *Rickettsiella* through lateral gene transfers from phylogenetically distant endosymbionts also inhabiting arthropods, like *Rickettsia* and *Wolbachia*. Both are common in arthropods [107, 108], especially *Rickettsia* in ticks [4, 109] in which co-infections with *Rickettsiella* have been reported [4]. These co-infections create physical proximity between these endosymbionts and may have facilitated lateral DNA transfers [110, 111], which could explain the intertwined evolutionary history of *cifA* and *cifB* genes among endosymbionts of arthropods.

The phylogenetic diversity pattern strongly suggests that horizontal transfers between arthropod species are key drivers of *Rickettsiella* spread, dictating global incidence of infections across arthropod communities. Repeated horizontal transfers results to a widespread and global distribution in diverse arthropod hosts, but also to multiple acquisitions of different *Rickettsiella* strains in the same host species as in *I. ricinus* (this study) and the polar seabird tick *Ixodes uriae* [25]. The ability of *Rickettsiella* to switch from one host species to another may therefore explain why no COGs specific to tick-borne *Rickettsiella* have been identified. There are, however, some exceptions: i) one uncharacterised protein is shared by all *Rickettsiella* isolated from *I. ricinus*, ii) the fatty acid oxidation complex subunit alpha is shared by all strains isolated from *I. ricinus* apart from *R. massiliensis* 20 B, iii) five clusters of hypothetical proteins specific to the Oerr strain, and nine clusters of uncharacterised proteins specific to the Opha strain. While most of these proteins have not been associated with any function yet, the fatty acid oxidation complex is quite interesting. Indeed, bacteria may exploit fatty acids from the host and use them for their own sustainment [112]. On the other hand, the oxidation of fatty acids might allow *Rickettsiella* to produce energy-rich compounds that could be used by *Rickettsiella* itself but also its tick host, aiding the host’s metabolism, as previously suggested for the tick microbiome [113, 114] and *Coxiella*-like endosymbionts [41, 115].

## Conclusions

*Rickettsiella* is a diverse group of endosymbiotic bacteria that exhibit a range of adaptations, from pathogenic to mutualistic, enabling them to thrive within arthropod cells and manipulate their hosts’ biology. Over the past decade, significant progress has been made in understanding their impact on arthropods and introducing these bacteria into the field of genomics. Our study highlights that *Rickettsiella* strains share similar metabolic capabilities and collectively lack virulence genes, suggesting a functional role in the arthropod hosts. In addition, we identified homologs of *cif* genes, associated with cytoplasmic incompatibility in Wolbachia, in two Rickettsiella strains, suggesting that they may use reproductive manipulation to spread in host populations (as previously shown for another Rickettsiella strain isolated from spiders). However, *Rickettsiella* remains an understudied aspect of tick biology, with tick-associated strains not forming a monophyletic clade, and a comprehensive approach is needed to fully understand its interactions with various arthropod hosts. Key questions remain, such as: How are the abundance, diversity, and distribution of *Rickettsiella* maintained worldwide? Can *Rickettsiella* be used effectively in pest control? How do they manipulate host reproduction? What roles do they play in the evolution of their arthropod hosts? Importantly, future research is needed to further investigate how *Rickettsiella* affects the fitness of their arthropod hosts, including ticks, and to what extent they can cause diseases in humans.

## List of abbreviations

*Ca.*: *Candidatus*
CI: Cytoplasmic incompatibility
*R.*: *Rickettsiella*
TCA: Tricarboxylic acid (cycle)
TIVASS: Type IV A secretion system
min: minutes
nt: Nucleotides
ANI: Average Nucleotide Identity
CDSs: Coding sequences
COGs: Clusters of Orthologous Genes
aa: Amino acids

## Declarations

The authors have complied with all ethical standards required for conducting this research. Consents and approvals are not applicable to this research. The genomes are available on ENA under project number PRJEB70514 (Ird1 and Ird6) and PRJEB82506 (Iric, Oerr and Opha). All the bioinformatics pipelines are available on GitHub (https://github.com/annamariafloriano/RickettsiellaComparative). This work was funded by French Agence Nationale de la Recherche (ANR, France, ref. ANR-21-CE02-0002, Laboratoire d’Excellence CEBA, ref. ANR-10-LABX-25-01 and LabEx CeMEB, ref. ANR-10-LABX-04-01), and by the United States Department of Agriculture (USDA, USA, grant no. 2019-67015-28981; https://www.asf-nifnaf.org/).

## Acknowledgements

We thank Carlos João Quembo, Florian Taraveau and Maxime Duhayon for their contribution in the sample collection. This work was performed using the computing facilities of the CC LBBE/PRABI.

## Figure captions

Supplementary Figure 1: barplot of intact genes and pseudogenes detected by PseudoFinder in each organism of Dataset R. The height of each bar represents the number of intact genes (blue) and pseudogenes (magenta).

Supplementary Figure 2: Intersection plot of Dataset R produced with the R package UpSetR, where all intersections are presented. In the left side, the horizontal bar plot represents the number of orthogroups present in each organism of the dataset. In the top, the vertical bar plot represents the number of orthogroups that belong to specific intersections of organisms in the dataset. The height of each bar indicates how many orthogroups are in that particular intersection, i.e., shared by the combination of organisms indicated by the intersection matrix below. In the intersection matrix, on the bottom, each row represents an organism and each column represents a possible intersection of these organisms. Filled circles indicate that the organisms in that row are part of the intersection shown in the corresponding column. The lines connecting the filled circles indicate which organisms are included in each specific intersection.

Supplementary Figure 3: Phylogenies of the thiamine genes present in Oerr and not in the other Rickettsiella strains of Dataset R, in order: *thiD*, *thiDE*, *thiO*, *thiG*, *thiS*. Although it is not possible to clearly establish the evolutionary origin of these sequences, it is clear that they are of Gammaproteobacteria/Pseudomonadota origin, often found in Legionella and in two other Rickettsiella organisms (See also Supplementary Table 3).

Supplementary Table 1: Metadata, genome statistics and number of intact and pseudo-genes of the organisms included in Dataset R. The ANI values for Dataset R and for the redundant genomes that were excluded from the analyses are also presented, as well as the mobile elements for each genome in Dataset R.

Supplementary Table 2: Orthogroups.GeneCount.tsv contains the orthologous clusters matrix produced with OrthoFinder on the translated intact genes of Dataset R and on the translated gene sequences of the outgroup B. aphidicola. This matrix was used to select the clusters for the phylogenomics-based tree and for the intersection plots. In the following sheets, the intersections of clusters of specific organisms of the datasets are presented, together with their KEGG and Prokka annotations for one of the organisms in the intersection (sequence ID reported). SharedByAll: clusters of intact genes shared by all organisms in the dataset; SharedByAllRickettsiella: shared by all *Rickettsiella* strains of Dataset R; SharedAllIricinus: shared by all *Rickettsiella* strains isolated from *I. ricinus* included in Dataset R; SharedAllIricinus_NoRmassiliensis: as SharedAllIricinus, but not present in the putative pathogen *R. massiliensis* 20B; Shared_MostPSstrains_IricRmassiliensis: shared exclusively by the *Rickettsiella* strains with the highest number of pseudogenes, i.e. *R. massiliensis* 20B and *Rickettsiella* strain Iric; SpSpecific_RDG: clusters exclusively present in the genome of the *Rickettsiella* endosymbiont of *D. gallinae*; SpSpecific_Rgrylli: clusters exclusively present in the genome of R. grylli; SpSpecific_Ird1: exclusively present in the genome of *Rickettsiella* strain Ird1; SpSpecific_Oerr: exclusively present in the genome of *Rickettsiella* strain Oerr; SpSpecific_Opha: exclusively present in the genome of *Rickettsiella* strain Opha; SpSpecific_Rmassiliensis: exclusively present in the genome of *R. massiliensis* 20B; SpSpecific_Rviridis: exclusively present in the genome of *R. viridis*.

Supplementary Table 3: KEGG annotation of the organisms of Dataset R obtained through BlastKoala (default settings). The column Knumbers indicates the KEGG ID. The values indicate presence (2), pseudogenization (1), and absence (0) in each organism. The sheets Summary NutrMutPaths, NutrMutPaths and EnergyPaths are focused on specific metabolic pathways potentially involved in nutritional mutualisms (B vitamins and heme metabolism, energy production); in these tables, “B”s mean null values in correspondence of the pathway names and were inserted as information for the script to obtain the heatmap shown in Figure 1. Thiamine_Oerr contains the amino acid sequences of the thiamine pathway components identified in *Rickettsiella* strain Oerr and not in the other *Rickettsiella* strains included in Dataset R, together with the five best-hits of blast against the non-redundant NCBI database, ordered by identity percentage (ID of the hit sequences, identity percentage, query cover, E value and available taxonomic annotation of the hit organism reported). The organisms and their sequence accession number used to infer the phylogenies of the Oerr thiamine genes (Supplementary Figure 3) are reported in the sheet Thiamine Phylogenies datasets.

Supplementary Table 4: Accession number of the *Rickettsiella*, *Wolbachia*, and *Rickettsia* genomes from public databases used to screen the cifA and cifB genes pair homologs.

Supplementary Table 5: Intersections of clusters of all encoded sequences, all intact sequences, and all pseudogenes of specific phylogenetic clades represented in Figure 1. A reference sequence ID is provided for each cluster, together with the KEGG (obtained with BlastKoala, default settings) and Prokka annotation.

Supplementary Table 6: Annotation of putative virulence factors in the organisms of Dataset R, annotated using amino acid Legionella queries downloaded from VFDB as a reference in a local blast search (default parameters) against translated all genes of the organisms of Dataset R. Sequence IDs are reported for each organism, cells are highlighted in yellow when the sequence was identified as a pseudogene by PseudoFinder. The hyphen (“-”) indicates complete absence of the sequence.

## Notes

### Competing Interest Statement

The authors have declared no competing interest.

### Summary of Updates

The article has been recommended by PCI Evolutionary Biology and we are providing the updated version.

