## Supplementary figures and images for "Comparative genomics of *Rickettsiella* bacteria reveal variable metabolic pathways potentially involved in symbiotic interactions with arthropods"

### Supplementary Figure 1

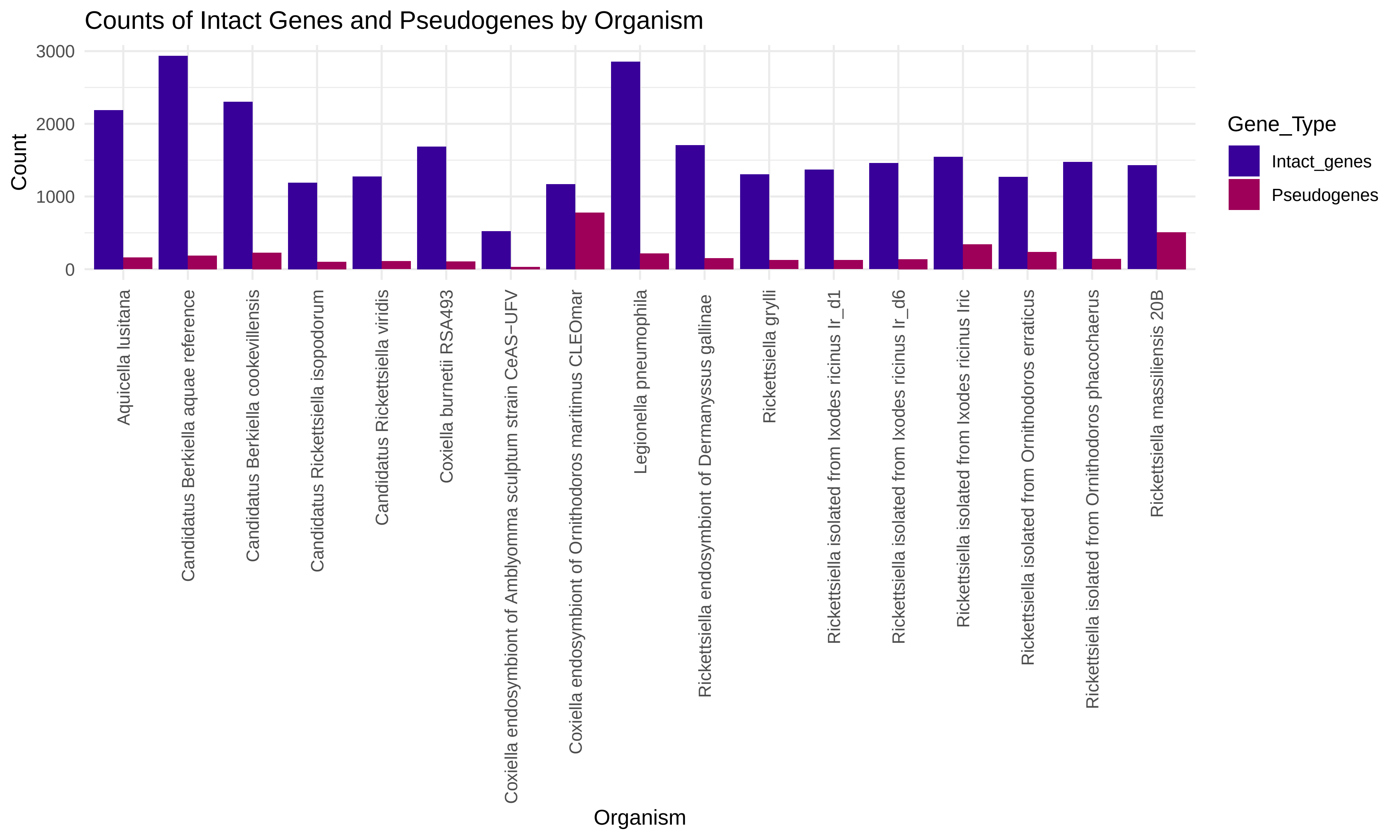

### Supplementary Figure 2

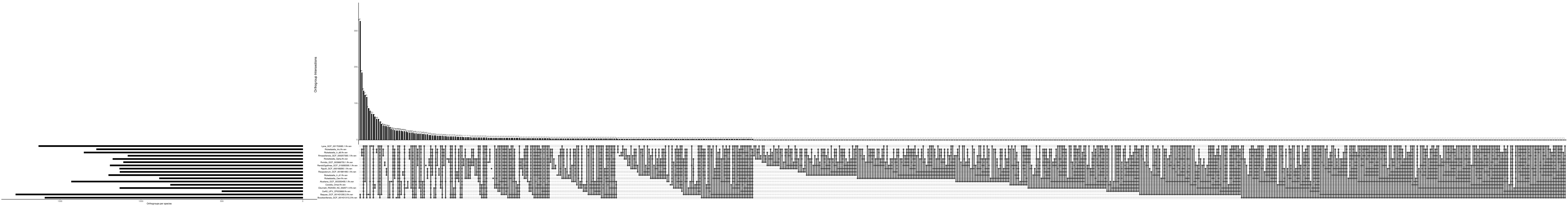

### Supplementary Figure 3

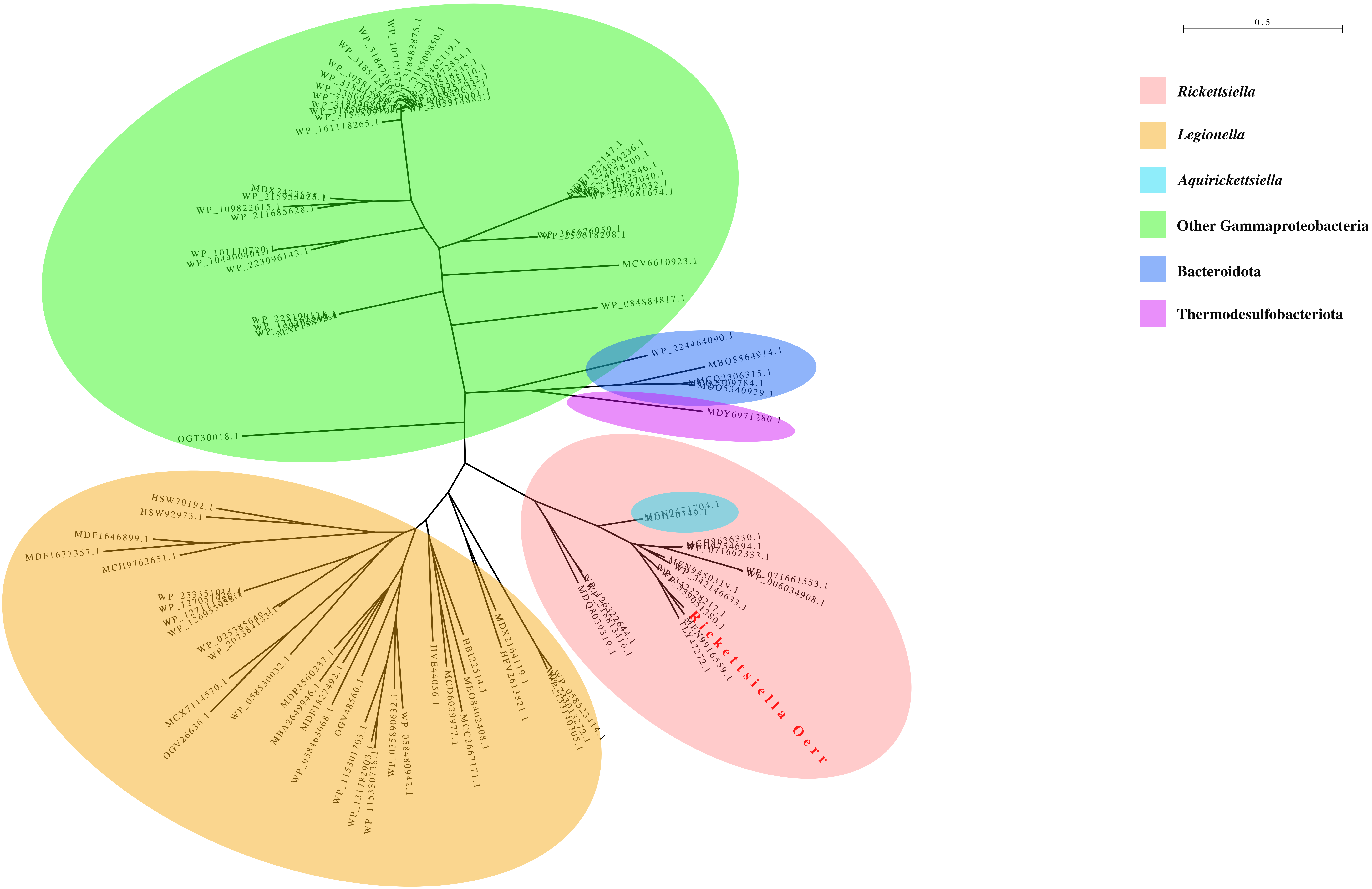

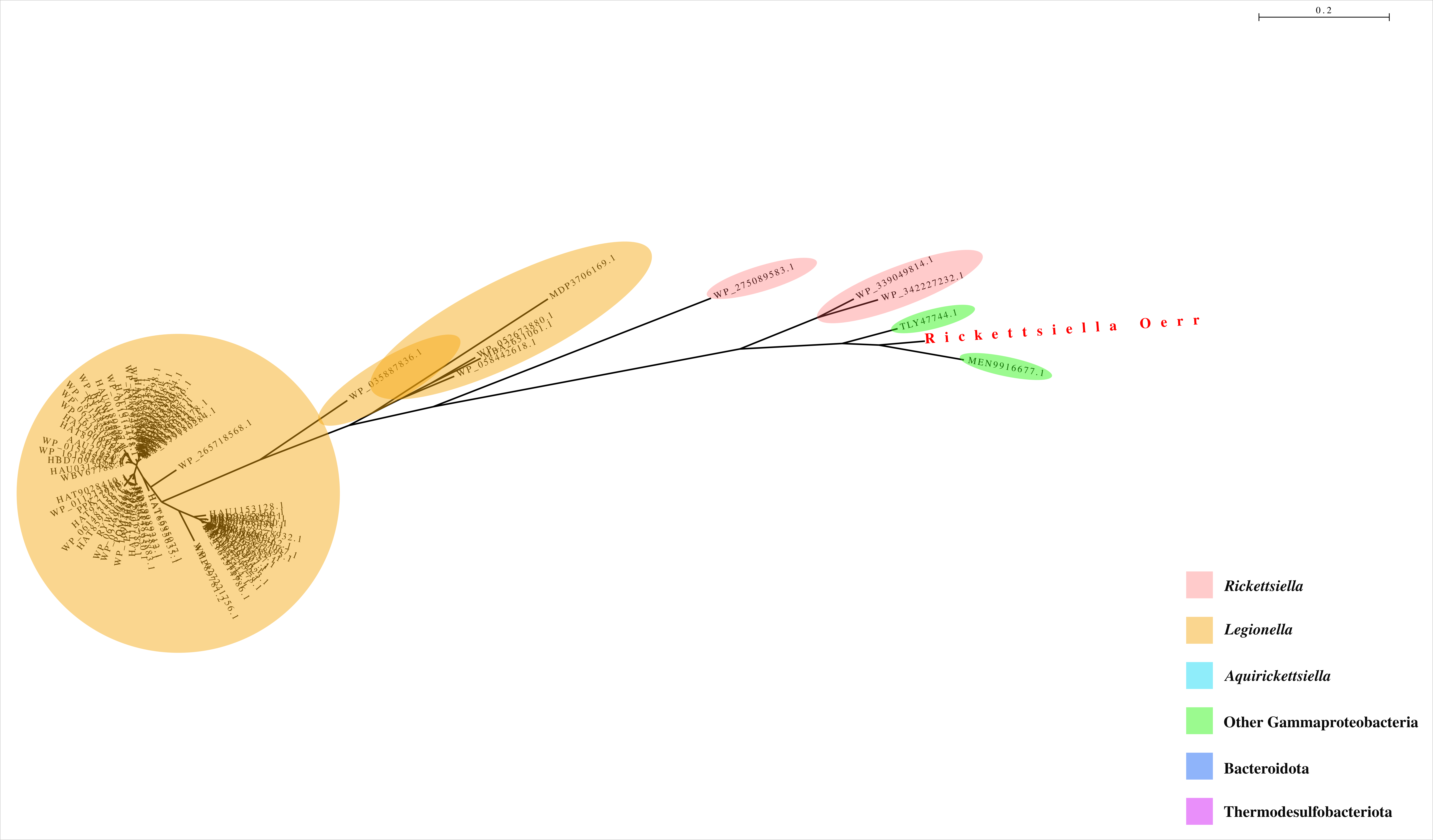

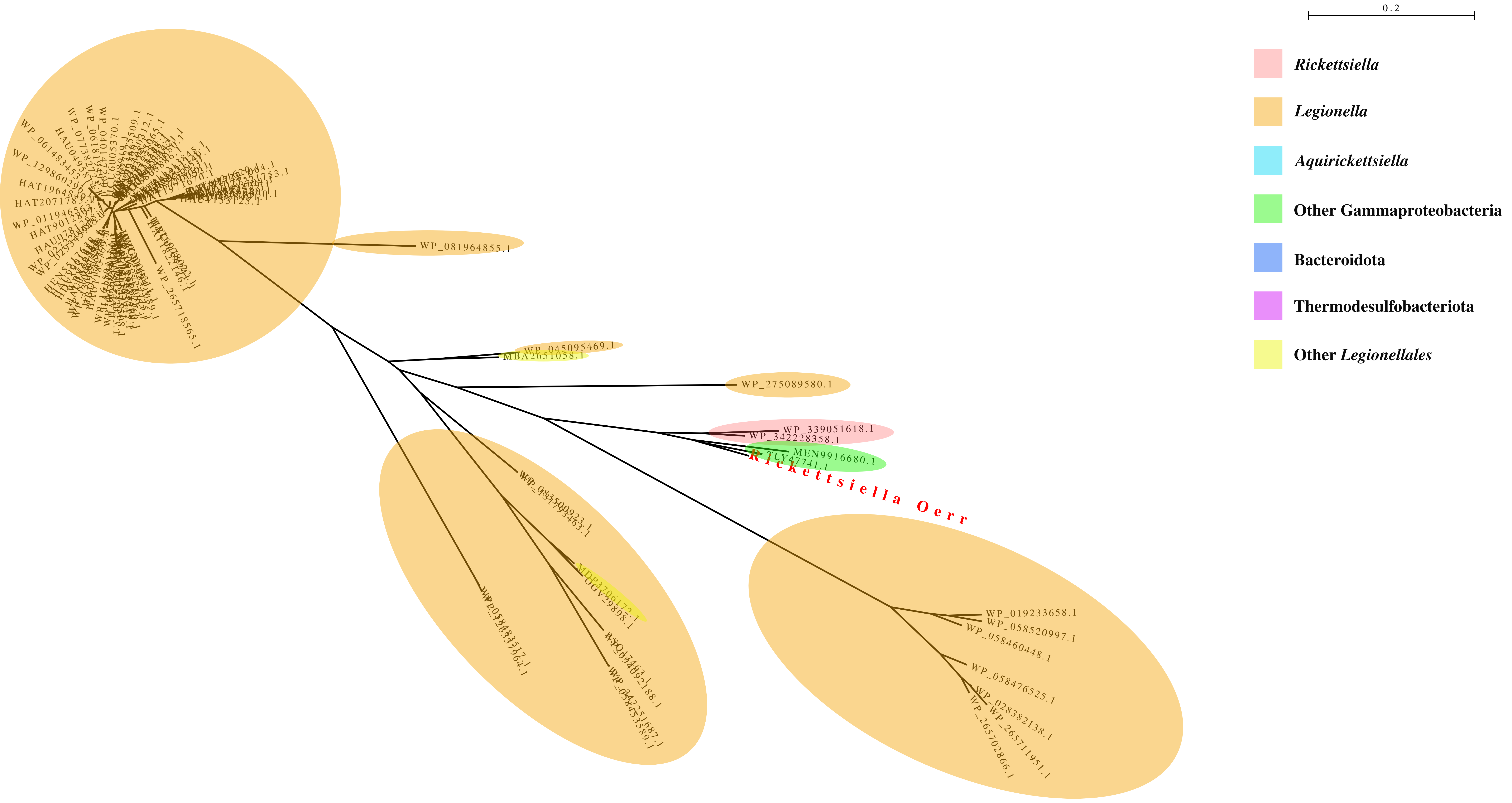

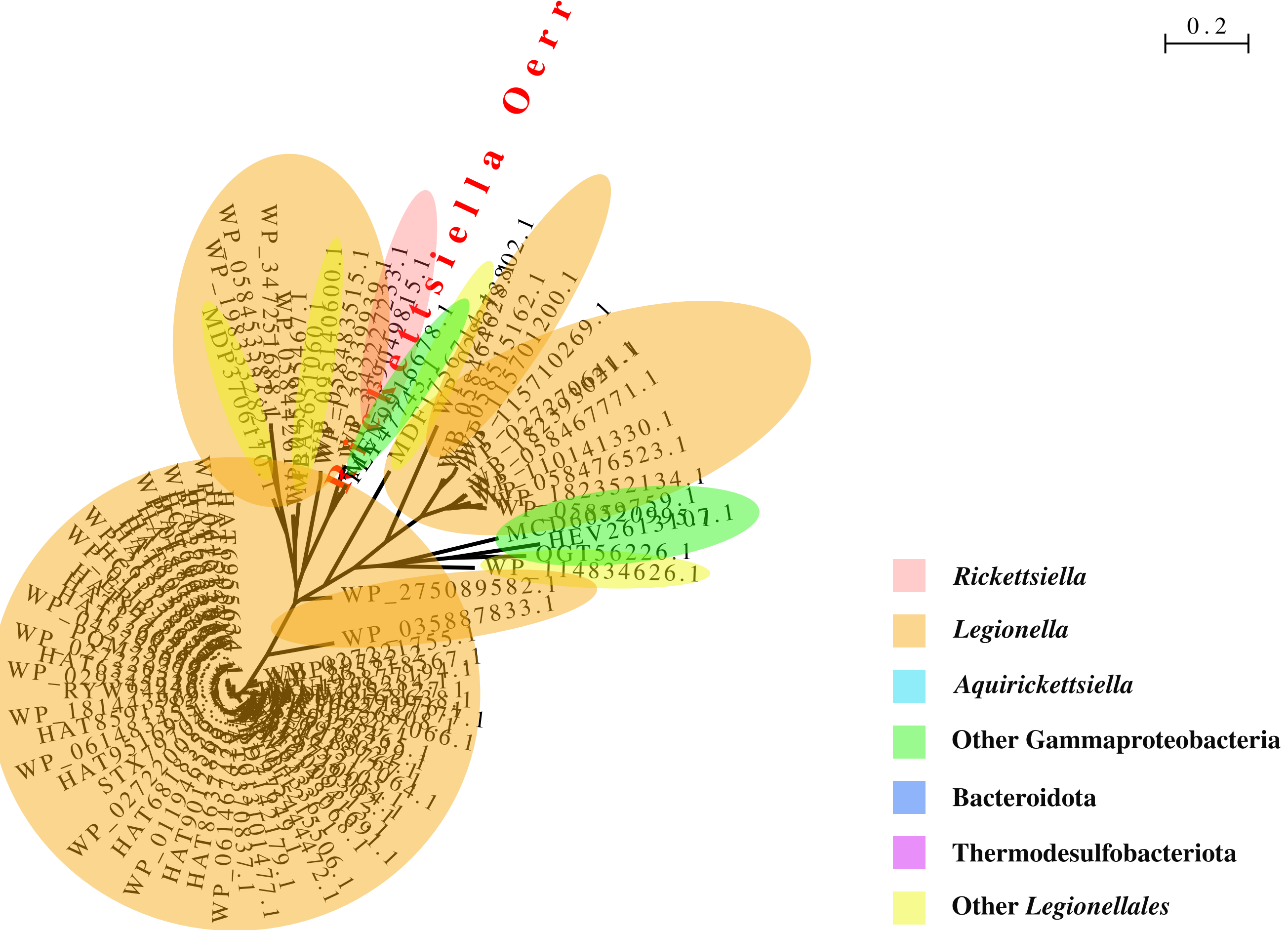

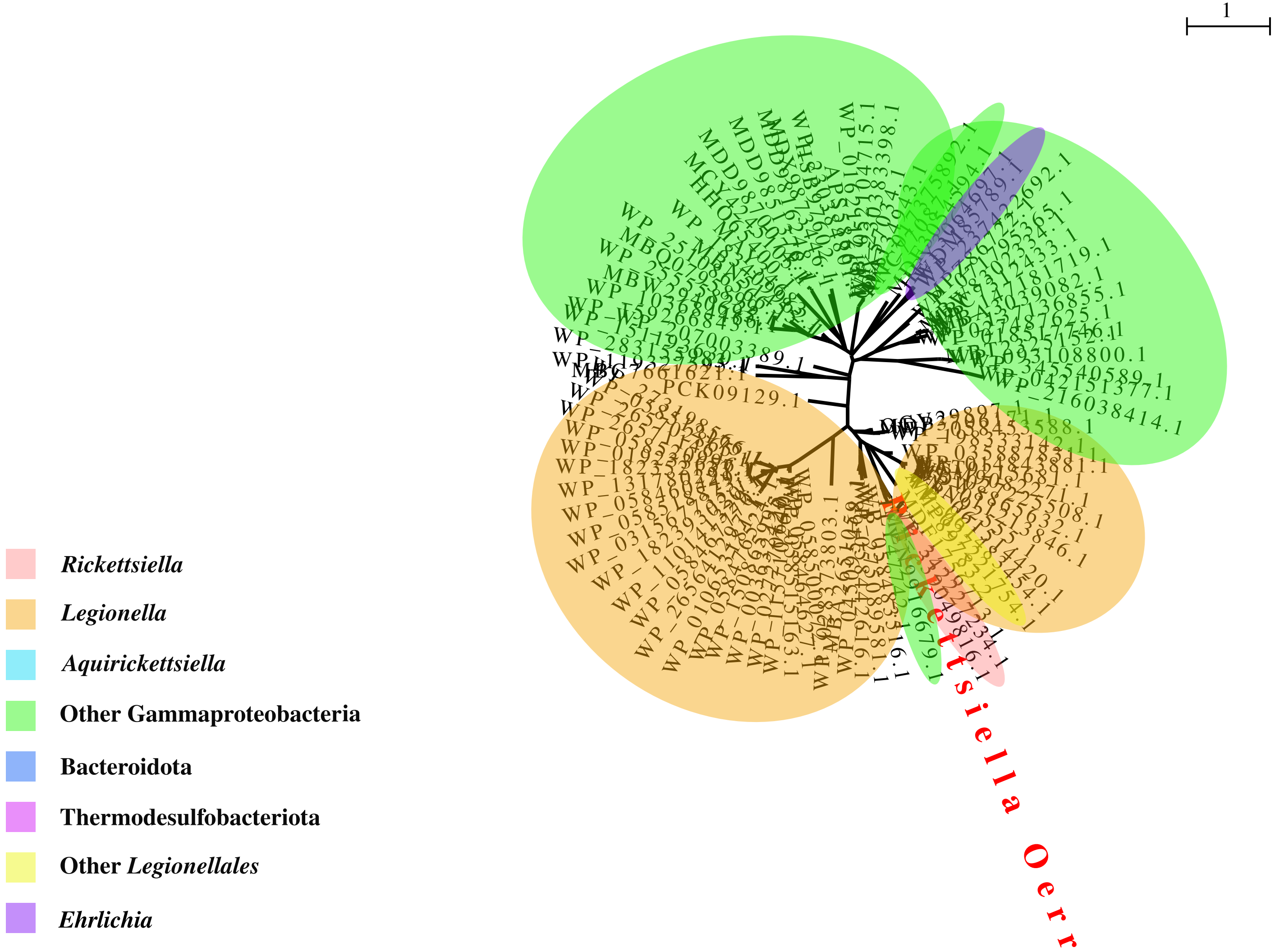
